# A repetitive mutation and selection system for bacterial evolution to increase the specific affinity to pancreatic cancer cells

**DOI:** 10.1101/217893

**Authors:** Masaki Osawa

## Abstract

It is difficult to target and kill cancer cells. One possible approach is to mutate bacteria to enhance their binding to cancer cells. In this study, Gram-negative bacteria *Escherichia coli* and Gram-positive bacteria *Bacillus subtilis* were randomly mutated, and then were positively and negatively selected for binding cancer and normal cells. With repetitive mutation and selection both bacteria successfully evolved to increase affinity to the pancreatic cancer cell line (Mia PaCa-2) but not normal cells (HPDE: immortalized human pancreatic ductal epithelial cell line). The mutant *E. coli* and *B. subtilis* strains could bind to Mia PaCa-2 cells about 10 and 25 times more than to HPDE cells. The selected *E. coli*, strain had mutations in biofilm-related genes and the regulatory region for a type I pilus gene. Consistent with type I pili involvement, mannose could inhibit the binding to cells. However, the results suggest that weak but specific binding is involved in the initial step of adhesion. To kill Mia PaCa-2 cells, hemolysin was expressed in the mutant strain. The hemolysin that was released from the mutant strain could kill Mia PaCa-2 cells. This type of mutation/selection strategy may be applicable to other combinations of cancer cells and bacterial species.

## Introduction

The main difficulty of cancer treatment attempting to target the cancer cells in the body, is that tumor specific surface markers are limited ^1, 2^. The surface of cancer cells is actually considerably different from normal tissue cells. However, almost all differences including protein expression, glycosylation, and lipid composition are quantitative; the difference is not the species of molecules but their amounts ^3, 4, 5^. It is possible that we may specifically tag cancer cells to attack them if we can develop an appropriate tool. One possible approach would be a probe that interacted weakly with multiple types of molecules that are expressed more highly on cancer cells. The cooperativity of multiple weak interactions could result in increased specificity for binding to cancer cells.

Bacteria may be a good candidate for this type of tool because of their large surface area and many surface ligands including protein, sugar moiety and lipid are available to develop interactions with cancer cells. One interesting observation is that various *B. subtilis* strains have largely different affinities for mucin, matrigel and a heterogeneous human epithelial colorectal adenocarcinoma cells line (Caco-2 cells) ^6^, suggesting that random mutations can affect the bacterial surface and determine the place where bacteria bind. Therefore it may be expected that a simple mutation/selection system may create bacteria that have higher affinity to cancer cells.

There are several advantages to use bacteria to fight cancer as follows. First, some bacteria have a natural capability to target cancer regions. The obligatory anaerobic bacteria such as *Bifidobacteria sp.* and *Clostridia sp.* prefer living in environments where there are no free oxygen molecules. Therefore we may expect that the hypoxic regions in tumors are a good home for them and in fact *C. novyi* was found to concentrate throughout cancer regions ^7^. This simple story, however, may not be everything because facultative anaerobic bacteria such as *E. coli* and *Salmonella*, which can live in both normal and hypoxic condition, can also accumulate in cancer regions in mice 8,9. There are some unknown mechanisms in cancer microenvironments which attract and/or support these bacterial growths.

Second, some bacteria naturally induce immunostimulation and attenuate cancer growth. One successful bacterial therapy for cancer has been to use BCG (Bacillus Calmette–Guérin) which is attenuated *Mycobacterium bovis.* Although the mechanisms are not fully understood, the main effects of BCG are to activate multiple immune pathways ^10^. In the case of *E. coli*, not only the tumor was cleared in a mouse model through CD8(+) T cells, but also in re-challenging mice that had cleared tumors, new tumors were rejected through both memory CD8(+) and CD4(+) T cells ^11^. Although these types of immunostimulation may be generalized with bacterial applications to various types of cancers, it is probably difficult and complicated for cancer treatments.

Third, bacteria can also work as a vehicle for drug delivery. One approach is to enhance the immunostimulation by expressing and secreting artificial cytokines from engineered bacteria ^12^. The simplest approach would be to release toxic molecules from bacteria to directly kill cancer cells. Secretion of a bacterial toxin such as alpha-hemolysin (aHL) and azurin have been shown to kill cancer cells in mice ^13-15^. Bacteria can also deliver converting enzyme for prodrug and show reduction of cancer mass in mice ^16,17^.

Pancreatic ductal adenocarcinoma is one of the deadliest cancers, with 4% survival rate for 5 years. It is very difficult to target pancreatic ductal adenocarcinoma cells. There is not even an effective biomarker for pancreatic ductal adenocarcinoma for diagnosis. However, one interesting approach has been selection of random aptamers reported by cyclic negative and positive selection for binding to secreted materials from non-cancer pancreatic epithelial cells vs pancreatic ductal adenocarcinoma cells. I have used a similar strategy in the present study. Instead of aptamers I have randomly mutated bacteria and applied negative and positive selection for binding to normal and cancer cells. The repetitive mutation/selection system produced bacteria which specifically bind to the pancreatic ductal adenocarcinoma cells.

## Methods

### Cells and bacterial strains

Immortalized human pancreatic ductal epithelial (HPDE) cells ^18^ and pancreas ductal adenocarcinoma cell lines (MIA PaCa2 cells) were kindly donated by Dr. M. S. Tsao, Ontario Cancer Institute and Dr. R. R. White, University of California San Diego. HPDE cells were cultured in Keratinocyte SFM with supplements and MIA PaCa2 cells were cultured in DMEM with 10% serum, unless otherwise specified. *E. coli* strain W3110 and *B. subtilis* strain 168 C were kindly donated by Dr. M. J. Kuhen, Duke University and Dr. J. Errington, Newcastel University, respectively. These bacteria were cultured in LB medium.

### Mutagenesis with UV irradiation

One ml samples of stationary cultured bacteria were frozen for stocks and thawed in 10 ml of LB. The suspension of bacteria was cultured at 37 C for 2 hrs for *E. coli* and 3 hrs for *B. subtilis*. One ml of bacteria was span down and resuspended in 0.5 ml of 0.1 M MgSO_4_. The bacteria were placed on a plastic dish as 20 µl X 25 drops and exposed to UV until 99% of bacteria were killed. This has been reported to cover all possible single mutations in genome ^19^. UV treated bacteria were cultured with 5 ml LB overnight and stored as frozen stock.

### Negative and positive selection for mutated bacteria against cancer cells

One ml of mutated bacteria was diluted 10 times in LB and cultured at 37 °C for 2.5 hr for *E. coli* and 3.5 hr for *B. subtilis*. Typically 100 µl of bacteria was spun down and resuspended with conditioned media of HPDE cells. HPDE cells were cultured as a confluent monolayer in T75 culture flasks (Corning) and just before addition of bacteria for negative selection, the medium was removed except for 1 ml left in the flask. The resuspended bacteria (100 µl) were added to the HPDE cells. To spread bacteria on the entire cell surface, the flask was tilted several times and incubated at room temperature for 20 min. During incubation the flask was tilted a couple of times to keep an even distribution of bacteria. The 1.1 ml supernatant, which contained the bacteria not bound to HPDE cells, was removed and transferred to positive selection system (Fig. 1).

**Fig 1.**
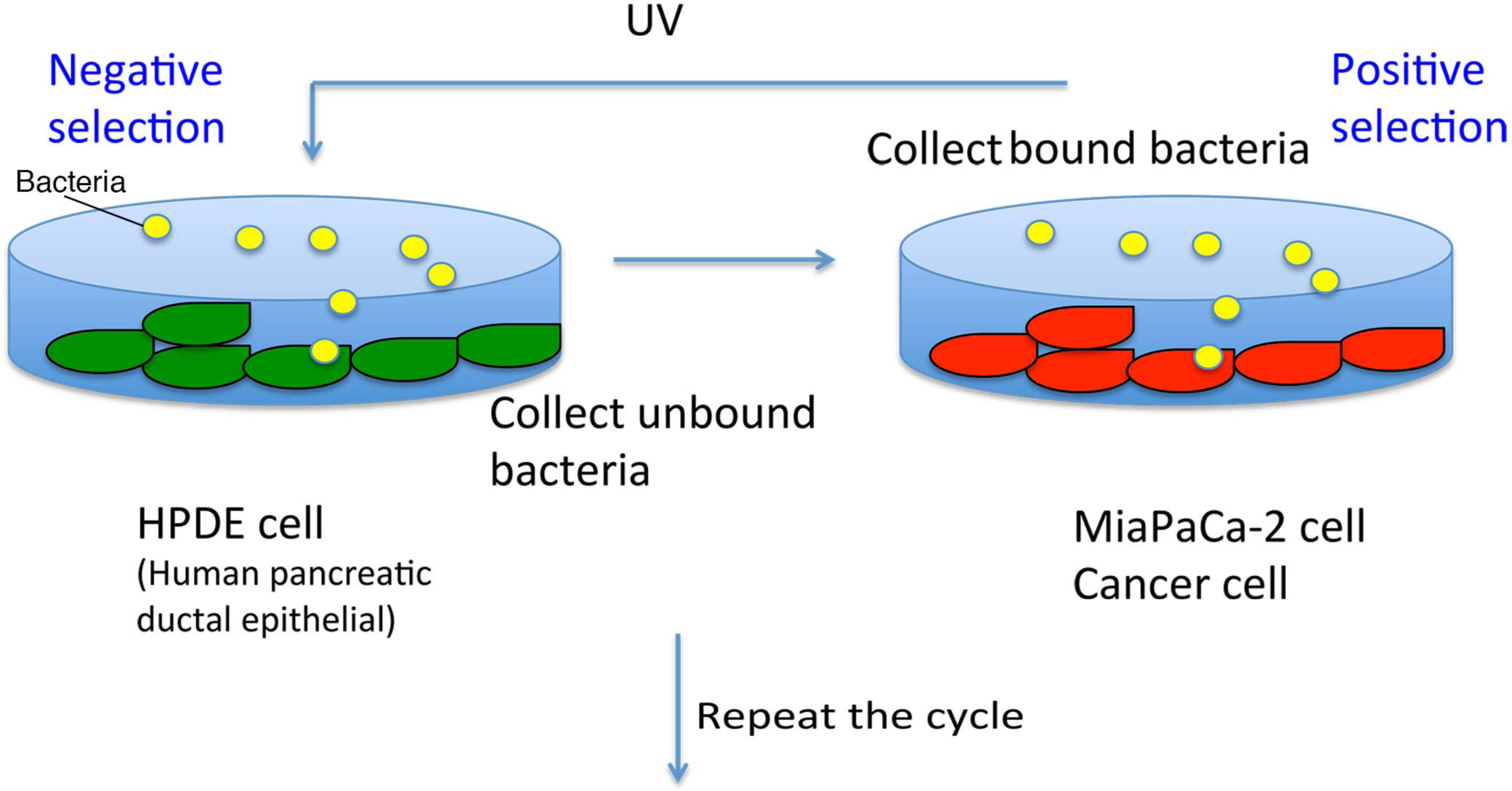
Repetitive mutaion and selection system.

For positive selection, MIA PaCa-2 cells were cultured to confluence in a T 25 flask. The medium was removed except for 1 ml left in the flask. The negatively selected bacteria were added to the MIA PaCa-2 cells and incubated for 10 min with occasional tilting to spread the bacteria. To remove bacteria that were not bound to the MIA PaCa-2 cells, the supernatant was removed by aspirator and MIA PaCa-2 cells were washed 10 times with 10 ml of PBS (Fig. 1).

The bacteria not only bind to the confluently cultured MIA PaCa-2 cells but also bind to the plastic surface exposed at the sides and top of the flask. To kill the bacteria stuck to the plastic surface, after carefully removing the PBS the flask was placed upside down and 1 ml of 70% ethanol was added. Because the flask was upside down, the ethanol spread on the top surface of the flask and killed bacteria attached there. The ethanol was replaced with a fresh 1 ml of 70% ethanol, and the flask was tilted 90 degree in 3 directions for 1 s and each time quickly put back to the upside down position. This process killed bacteria stuck on the 3 sidewalls. With the flask still upside down, the ethanol was removed and the flask neck and exit were wiped with a swab soaked with 70% ethanol to killed the bacteria stuck there. Then 5 ml LB was added and flask was flipped upright. The bacteria stuck on the cells were cultured overnight then stored at −80 C. The frozen stocks were thawed and used for the next cycle of radiation.

### Bacterial adhesion assay

The bacteria were grown in LB overnight and diluted 10 times with LB. Then the bacteria were cultured at 37 °C for 2 h for *E. coli* and 3 h for *B. subtilis*. One ml of bacteria was spun down and resuspended to 0.1 ml PBS with 10 µg/ml FM4-64FX, a red fluorescent dye that stains the bacterial membranes. After 25 min incubation the bacteria were washed twice with 1ml PBS and resuspended in 1 ml conditioned medium from MIA PaCa-2 or HPDE cells. Fifty µl (for *E. coli*) or 150 µl (for *B. subtilis*) of the bacterial suspension (containing 0.3 - 0.7 × 10^8^ bacteria as assayed by colony formation) was diluted with 0.95 (for *E.coli*) or 0.85 ml (for *B. subtilis*) of conditioned medium and the adhesion assay was started by replacing culture medium of the cells with the diluted bacterial suspension. After 10 min incubation, the cells were washed with 1 ml PBS three times and subsequently fixed with 4% formaldehyde for 10 min. The fixed cells were washed 4 times with 1 ml PBS and stored at 4 C.

The cultured cells were sparse for the adhesion assays. In some cases both MIA PaCa-2 and HPDE cells were cultured under identical conditions to test whether culture conditions affected the adhesion assays. In this case cells were plated on a glass surface coated with a fibronectin fragment FN7-10 containing the RGD peptide ^20^, and grown in Keratinocyte SFM with supplements.

### aHL expression and killing Mia PaCa-2 cell

The aHL gene was purchased as a double strand DNA fragment (gblock, integrated DNA Technologies) with flanking EcoRI – HindIII sites. The aHL gene was spliced into three kinds of pBAD based plasmids for expression in *E. coli*. The first construct was pBAD18 ^21^, which was referred to as pBAD_18aHL. The second construct improved the Shin-Dalgano region of pBAD18_aHL. The sequence ctagcgaattc ATG (start codon) was replaced with ctaacaggaggaattaaccATG (start codon) by doing PCR for the entire plasmid followed by ligation. This plasmid was referred to as pBADMO_aHL, and is expected to give a higher expression level of aHL. To make the third plasmid, the araE gene with Pcp8, which is a constitutively active promoter, was obtained by PCR from the genomic DNA of BW27783 strain ^22^ and was inserted between the M13 intergenic region and pBR322 ori in pBADMO_aHL by creating AscI – NcoI site in both pBADMO_aHL and Pcp8-araE. This plasmid was referred to as pBADMOE_aHL, and is expected to give a more uniform level of expression of aHL from cell to cell ^22^.

The mutant *E. coli* strain ECUV10c (cloned mutant *E. coli* treated with UV irradiations/selections 10 cycles) was transformed with each of the three plasmid and single colonies were cultured overnight in LB. For aHL induction, the cultures were diluted 20 or 100 times in LB with 0.2% arabinose grown in a shaker at 37 °C for 4 h or overnight. After induction, *E.coli* was spun down and supernatant, containing aHL released into the medium, was collected. The aHL was concentrated 40 times by centrifugal filter with cut off 10 kDa (Sigma). This aHL was applied to Mia PaCa-2 cells cultured in bottom glass dishes (Mattech), after diluton to 560 times in Mia PaCa-2 culture media, giving a 14-fold dilution from that in the original culture supernatant.

### Microscopy

Differential interference contrast and fluorescence images of bacteria and cells were obtained with a Zeiss Axiophot microscope with 100x (NA 1.3) or 40x (NA 1.3) objective lens and a CCD camera (CoolSNAP HQ, Roper). The FM-64 signal was captured through a rhodamine filter cube (BP546 FT580 BP590).

## Results

### Repetitive mutation/selection system

I took a simple strategy to obtain bacterial mutants that specifically bind to cancer cells. Bacteria randomly mutated with UV irradiation were subjected to negative selection with immortalized human pancreatic ductal epithelial (HPDE) cells and then positive selection with a pancreas ductal adenocarcinoma cell lines (MIA PaCa2 cells) (Fig. 1 and see materials and methods). To concentrate bacteria that specifically bind to the cancer cells, 3 rounds of negative and positive selection without UV irradiation were applied after each first mutation/selection process (Fig. 1). In many cases a reduced number of bacteria was applied for these additional selections to effectively concentrate the bacteria that have affinity to cancer cells. In general, 10^8^ bacteria were input to the negative/positive selection after UV irradiation, then 10^7^, 10^6^, and 10^5^ bacteria were input for second, third, forth cycles of selection without UV irradiation. These numbers of bacteria should retain some genomic variation.

These complete selection processes were repeated 10 and 9 times for two bacterial species, Gram-negative *E. coli* and Gram-positive *B. subtilis*. I chose these bacteria because their surfaces are very different; the Gram-negative bacteria have an outer membrane, while Gram-positive bacteria do not. This should give general insights into the mechanisms by which bacteria might increase their affinity to cancer cells by repetitive mutation/selection.

### Selected *E. coli* could specifically bind to Mia Paca-2 cells

To evaluate the bacterial adhesion to cancer cells, selected strains of both *E. coli* and *B. subtilis* were stained with the membrane dye FM4-64 and were applied to cells for 10 min as shown in Fig. 2. The mutant *E. coli* (ECUV10c: cloned mutant *E. coli* treated with UV irradiations/selections 10 cycles) showed the highest attachment to Mia PaCa-2 cells, which was about 10 times more than to HPDE cells (Fig. 2). In contrast, the original wt *E. coli* gave the lowest attachment to both Mia PaCa-2 cells and HPDE cells (Fig. 2 e, f, g, h, i). These results suggest that this mutation/selection system actually functions and successfully selected the mutant *E. coli* that specifically bind to the cancer cells. Interestingly, the intermediate mutant strain (ECUV4: heterogeneous mutant *E. coli* treated with 4 cycles of mutation/selection) increased the adhesion to both HPDE and Mia PaCa-2 cells compared to wt *E. coli*). The further cycles of mutation/selection to ECUV10c decreased adhesion to HPDE and increased adhesion to Mia PaCa-2 (Fig. 2 i). These results suggest that both positive and negative selections are functioning well.

**Fig. 2.**
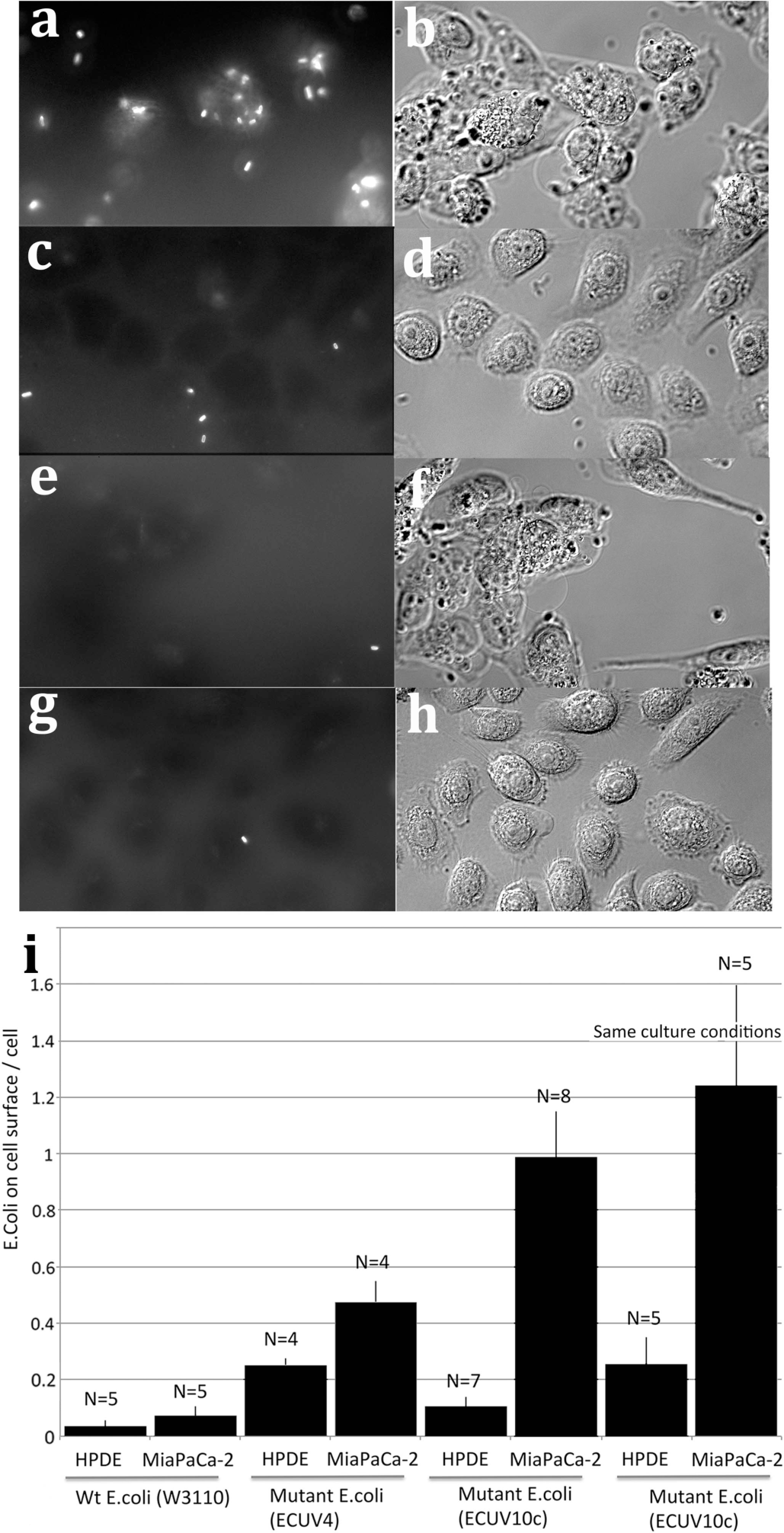
Mutant *E. coli* ECUV10c shows enhanced adhesion to pancreatic cancer cells. (a, b) Adhesion assay for ECUV10c with Mia PaCa-2 cells. (c, d) Adhesion assay for ECUV10c with HPDE cells. (e, f) Adhesion assay for wt *E. coli* with Mia PaCa-2 cells. (g, h) Adhesion assay for wt E. coli with HPDE cells. (a, c, e, g) show the fluorescently stained bacteria, and (b, d, f, h) show DIC images of the cells. (a, c) are superimposed images from two focal plane to show maximum number of *E. coli* on the surface of the same cells in each DIC images (b, d). (i) Quantification of the adhesion assays. Approximately 0.5 × 10^8^ bacteria were added to the cells. The exact number confirmed by colony assay was 0.3 – 0.7 × 10^8^. The number of bacterial was normalized to 0.5 × 10^8^. The number of attached bacteria adhered to all focal levels of the cells were counted and averaged per cell. The intermediate mutant strain ECUV4 showed increased adhesion to both HPDE and Mia PaCa-2 cells. The further selection to ECUV10c decreased adhesion to HPDE and increased adhesion to Mia PaCa-2 cells. The last two columns show adhesion to HPDE and Mia PaCa-2 cells plated on a glass surface coated with FN7-10 and grown in Keratinocyte SFM medium.

One concern was the different culture media used for HPDE cells and Mia PaCa-2 cells, which are cultured in Keratinocyte SFM with supplements, and DMEM/10% serum, respectively. To circumvent this difference, I attempted to culture Mia PaCa-2 cells in Keratinocyte SFM and found that they did not adhere to glass (or cell culture plastic) when they were plated. This may suggest that Mia PaCa-2 cells do not produce enough fibronectin by themselves. Therefore I coated the glass surface with fibronectin fragment FN7-10, which includes the RGD peptide for integrin binding ^20^. Mia PaCa-2 cells could adhere and spread on this substrate in Keratinocyte SFM and grow with very similar morphology to the Mia PaCa-2 cells in DMEM/10% serum. Under this condition, the ECUV10c could attach to the Mia Paca-2 cells at essentially the same level as the Mia Paca-2 cells cultured in DMEM/10% serum (Fig. 2 i). Adhesion to HPDE was increased somewhat, but the large differential was maintained under these identical culture conditions.

The complete genome of the mutant (ECUV10c) were determined by Illumina sequencing, and the differences from the database W3110 strain (accession # PRJNA16351) are shown in table 1. Our laboratory wild-type W3110 strain, the parent of ECUV10c), had 9 mutations from the database strain, including a mutation in fliI, the flagellum-specific ATP synthase. This is in agreement with the motility defect of lab strain W3110. This strain acquired 25 additional point mutations and an obvious large deletion during 10 cycles of UV irradiation with positive/negative selection, which created the mutant ECUV10c. The large deletion corresponds to the e14 which is a UV-excisable defective prophage with 19 genes and 3 pseudogenes, *ymfO'*, *ymfP'* and *stfE'*, located at 1197893 – 1212987 in w3110 genome in PRJNA16351. This is reasonable because this strain received UV irradiation. Eight of 25 mutations were in the icd (isocitrate dehydrogenase) gene right next to e14 deletion. Notably all these 8 mutations in icd are silent mutations.

Three of the mutations are in the genes involved in biofilm formation (*ycdR, ydeH, yjfO*). These may affect the binding of ECUV10c to cancer cells by changing the surface of the bacteria. Probably the most important mutations are in the Fim genes encoding type I pili, which have affinity to mannose residues. The mutations are located at 3 nucleotides before the transcription start of *fimB* gene, and in the ORF of the *fimE* gene. Both FimB and FimE are recombinases and regulate the expression level of type I pili by flipping the promoter region of FimA which is the axis of the type I pili.

### Involvement of Type I pili in *E. coli* adhesions

To check the involvement of type I pili for the binding of *E. coli* to the cancer cells, 1% mannose was added to the medium during the adhesion assay. The adhesions of both ECUV4 and ECUV10c were reduced to less than those of wild type, suggesting that the adhesions are highly dependendent on the type I pili (Fig. 3). However, these type I pili-dependent adhesions were probably regulated by additional factors. In the case of ECUV4, the type I pili-dependent adhesions for Mia PaCa-2 and HPDE cells were increased by the mutation/selection system (Fig. 2, Fig. 3). Because there are no FimB/E mutations in this ECUV4 population (table s1), some other factors should enhance these adhesions to both Mia PaCa-2 and HPDE cells through type I pili. The mutation in *ycdR* gene may be responsible for this adhesion enhancement because about half of population in ECUV4 has a mutation in this gene (see Supplementary Fig. S1a online).

**Fig 3.**
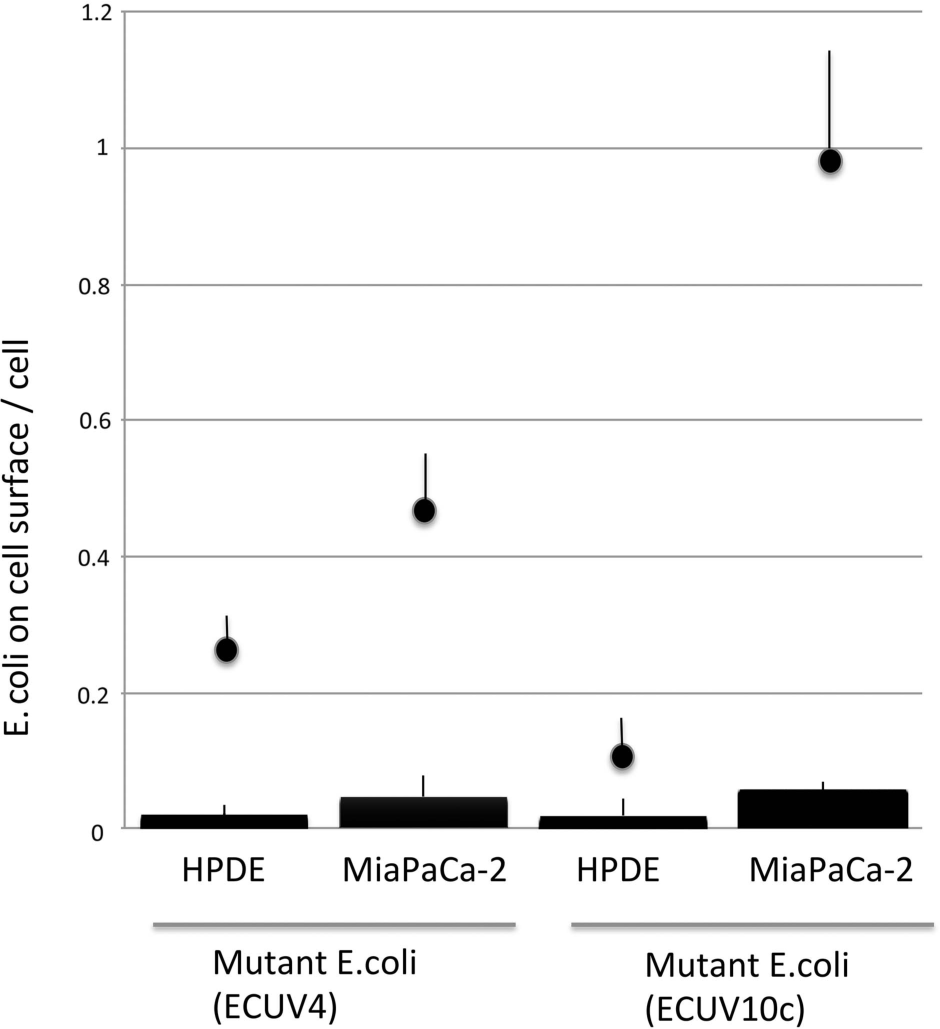
Mannose inhibition of mutant *E. coli* adhesion to cells. The adhesion assays were applied to ECUV4 and ECUV10c in the presence of 1% mannose. Adhesion in mannose is indicated by the bars at the bottom. Dots with error bars are replotted from Fig. 2 (without mannose).

The ECUV10c strain has a mutation in both *fimB* and *fimE* regions and probably has increased number of the type I pili, which may increase the type I pili-dependent adhesions on Mia PaCa-2 cells (Fig. 2). However, the same type I pili-dependent adhesions on HPDE cells were actually reduced for ECUV10c compared to ECUV4, again suggesting that there are some additional factors involved (see discussion and Fig. 7).

### Killing Mia PaCa-2 cells with alpha-hemolysin released from ECUV10c

In order to kill Mia PaCa-2 cells, *Staphylococcus aureus* aHL was expressed in ECUV10c, following reports that *E. coli* expressing aHL kills cancer cells both in cell culture and in mice ^13,14^. It is known that this method has difficulties because the sensitivity to aHL varies for different cancer cell lines, and also overexpression of aHL lyses bacteria ^13, 14, 23^. To address these problems, I constructed three arabinose-inducible pBAD-based plasmids, which should give different expression levels of aHL. The supernatant of the *E. coli* expression cultures was tested for aHL by adding to culture medium of Mia PaCa-2 cells. When aHL was expressed from pBAD18_aHL in ECUV10c, the supernatant did not kill Mia PaCa-2 cells.

To increase the expression level of aHL, I made a second plasmid, pBADMO_aHL, which has an improved ribosomal binding site from pBAD18_aHL. The supernatant medium from 4 h induction of ECUV10 with pBADMO_aHL could kill Mia PaCa-2 cells. The supernatant was concentrated 40 times with a centrifugal filter and diluted 560 times in the culture medium of Mia PaCa-2 cells, resulting in 14 fold dilution from the original supernatant.

To further enhance the expression level, a third plasmid, pBADMOE_aHL, was created. For this I incorporated into pBADMO_aHL an additional gene expressing AraE with a constitutively active promoter. It is known that expression of AraE (low affinity high capacity arabinose transporter) increases the protein expression level from pBAD, and also generates a more uniform response of individual bacteria. When aHL was expressed for 4 hrs from pBADMOE_aHL in ECUV10c, the supernatant medium, Mia PaCa-2 cells were killed at four times dilution from the concentration of the original supernatant. However, higher dilution did not have activity to kill Mia PaCa-2 cells, indicating that aHL concentration from pBADMOE_aHL is lower than that from pBADMO_aHL. The reason for this is that ECUV10c with pBADMOE_aHL did not grow well in induction media, probably because the higher and uniform expression of aHL lysed ECUV10c, consistent with the previous report (see Supplementary Fig. S2 online) ^13^. In contrast, ECUV10c with pBADMO_aHL could apparently grow while it released aHL into the media. As a result, the aHL activity (concentration) is higher in the media for ECUV10c with pBADMO_aHL than with pBADMOE_aHL.

Mia PaCa-2 cells remained attached to the glass surface 1 h after addition of supernatant of ECUV10c with pBADMO_aHL, but they started shrinking and some of them had distinctive bubble like structures (Fig. 4b). After 4 hr most cells were rounded up (Fig. 4c). After overnight treatment almost all cells were detached from glass surface (Fig. 4d, e).

**Fig 4.**
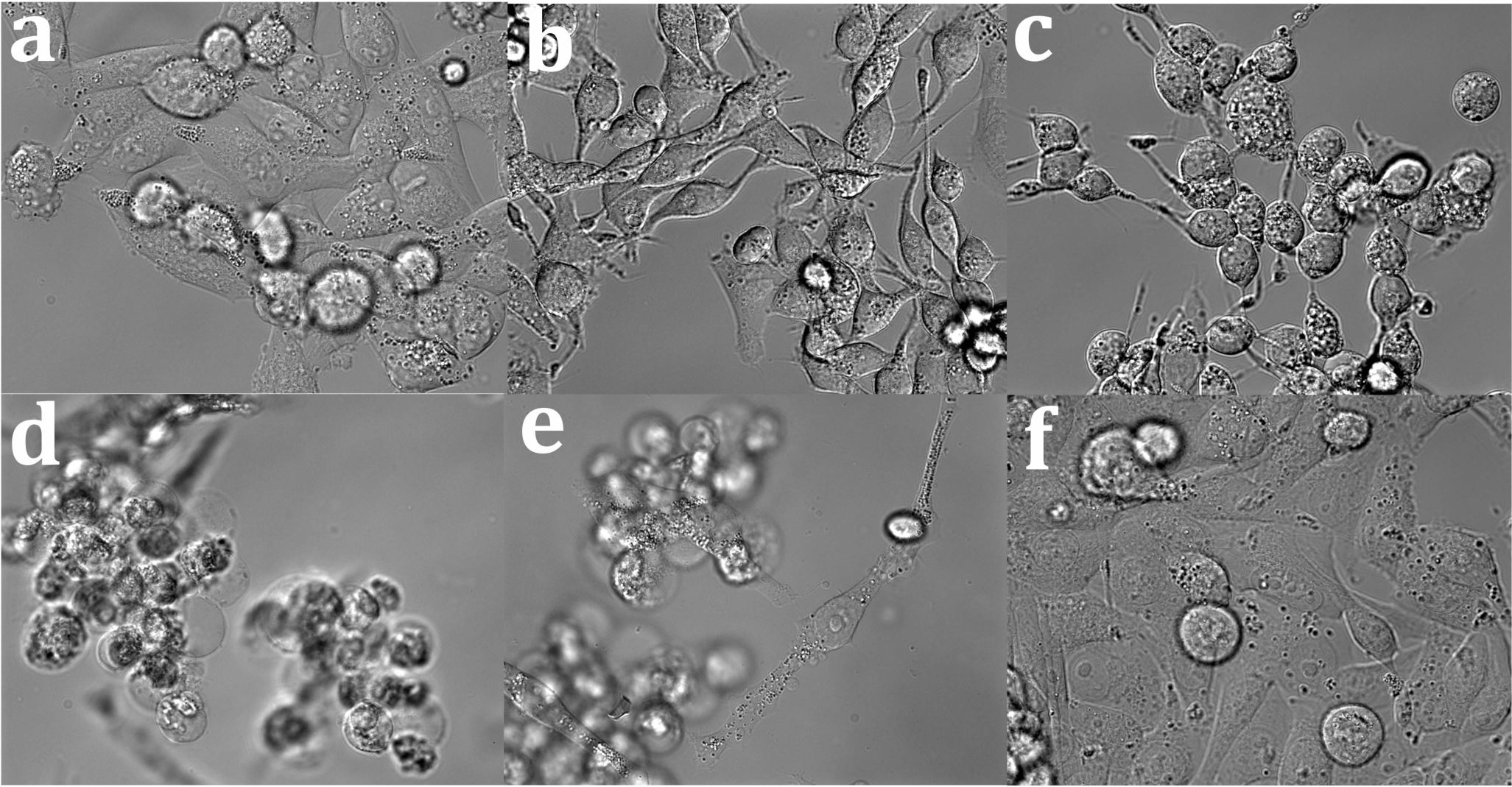
Killing Mia PaCa-2 cells by hemolysin released from ECUV10c. ECUV10c with pBADMO_aHL was induced for hemolysin expression with 0.2% arabinose. The supernatant of the culture was collected after 4h induction. 1.8 µl of concentrated supernatant was added to Mia PaCa-2 cells in 1ml of culture media, which corresponds to 14 times dilution from original ECUV10c supernatant. The images show cells (a) before addition of supernatant, (b) 1 h, (c) 4 h, and (d, e) overnight after addition. (f) is negative control with Mock plasmid.

### Selected *B. subtilis* could specifically bind to Mia Paca-2 cells

The same selection process and adhesion assays were applied to *B. subtilis*. FM4-64 could stain *B. subtilis* very well as shown in Fig. 5. Although some transfer of the dye from attached *B. subtilis* to the cell membrane after fixation was observed, *B. subtilis* on cells were easily detected by fluorescence and confirmed in DIC images. Unlike *E. coli*, many mutant *B. subtilis* were found beneath cells probably because of their active motility. BSUV9c, the cloned mutant *B. subtilis* with 9 cycles of UV irradiation/selection, attached to Mia PaCa-2 cells 25 times more than to HPDE cells (Fig. 5 a, b, c, d, i). In contrast, the original wt *B. subtilis* attached to HPDE cells at about the same level as BSUV9c. The mutations in this case resulted in a large increase in attachment to Mia PaCa-2.

**Fig 5.**
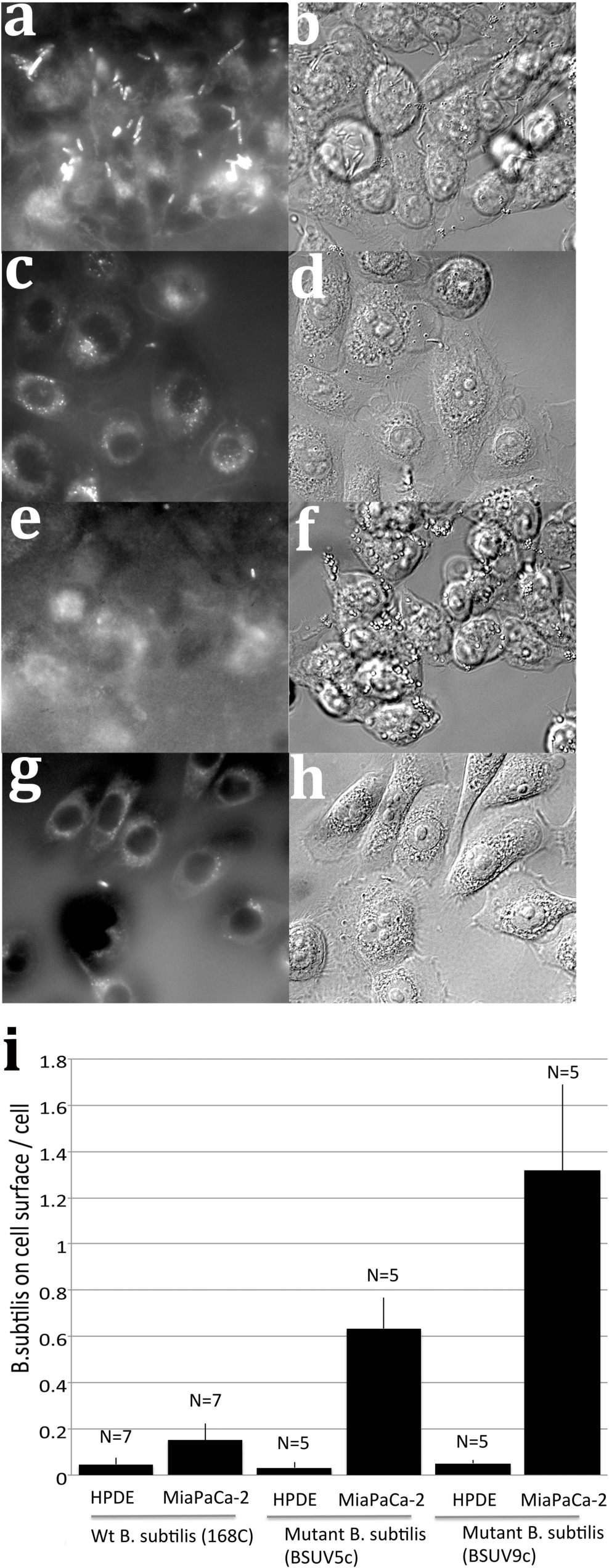
Increase of mutant *B. subtilis* adhesion to pancreatic cancer cells. (a, b) Adhesion assay for BSUV9c with Mia PaCa-2 cells. (c, d) Adhesion assay for BSUV9c with HPDE cells. (e, f) Adhesion assay for wt *B.subtilis* with Mia PaCa-2 cells. (g, h) Adhesion assay for wt *B. subtilis* with HPDE cells. Fluorescence images of FM4-64FX-stained bacteria are in (a, c, e, g) and DIC images of cells are in (b, d, f, h). (i) Quantification of the adhesion assays. The quantification was performed as described in Methods and Fig. 2

Because *B. subtilis* cells are highly motile, movies were captured to see how the interaction between cells and *B. subtilis* occurs (Supplementary movies SI1-4 show different combinations of bacteria and cells). Interestingly the initial attachments were formed at the leading pole of BSUV9c with the cell surface, implying that the molecules which has affinity to cell surface might be accumulated at the pole. The attachments were only occasionally seen for wt *B. subtilis* with both Mia PaCa-2 and HPDE cells.

In the case of mutant *B. subtilis*, the attachments to HPDE cells are as low as wt *B. subtilis*, whereas the attachments to Mia PaCa-2 cells are dramatically increased (see Supplementary movie SI1-4 online). This suggests that the initial attachments were also increased to cancer cells by mutation/selection system. These initial attachments on the cell surface were much more numerous than the bacteria per cell in the adhesion assay, suggesting that many of these bacteria were weakly attached and released by the washing process before fixation (Fig. 6 c, d).

**Fig 6.**
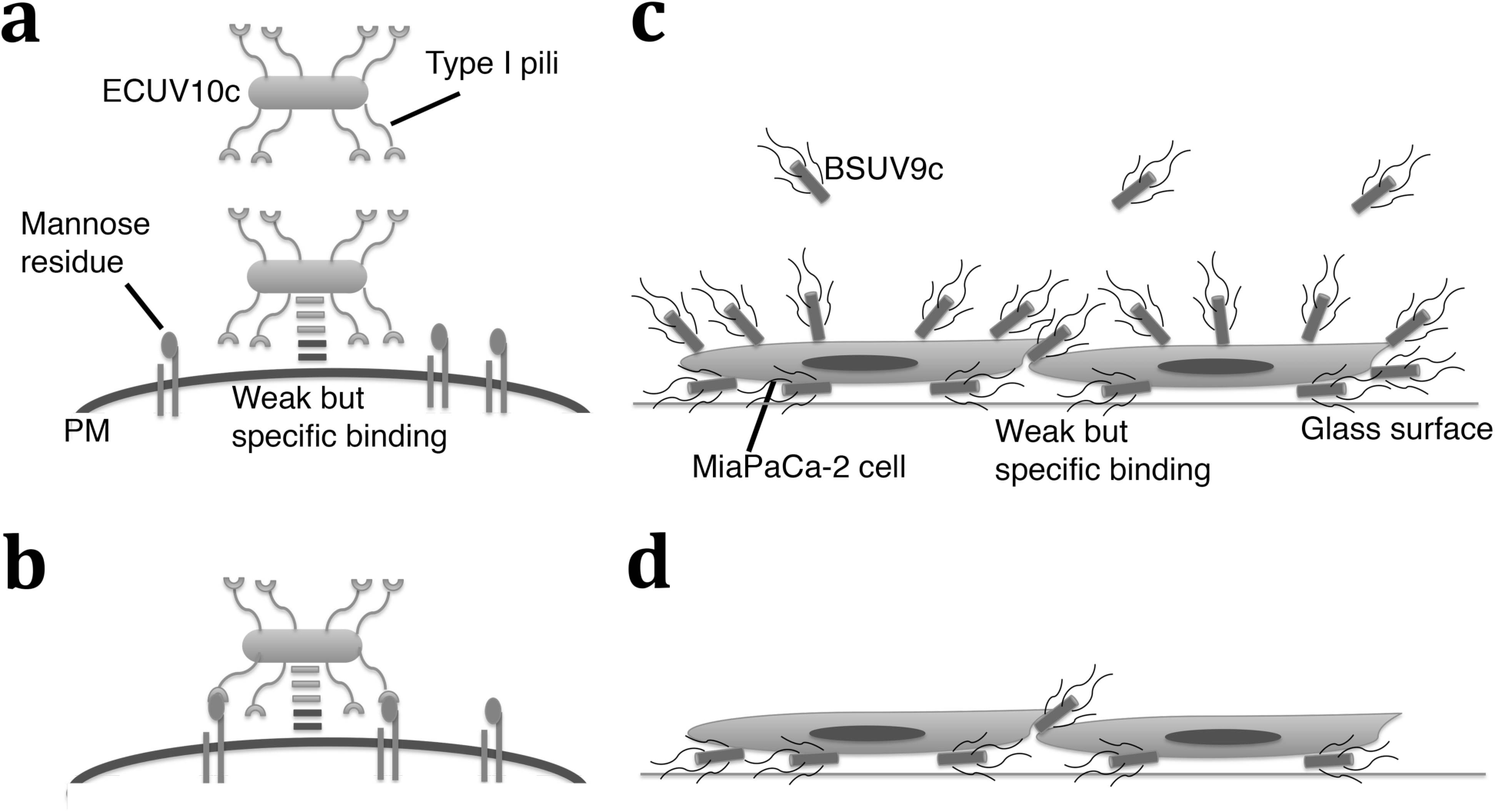
Models for how ECUV10c and BSUV9a may bind to the Mia PaCa-2 cells. (a) ECUV10c specifically binds to the Mia PaCa-2 cell surface through an unknown mechanism. Because of the low affinity of the binding, ECUV10c can be detached from Mia PaCa-2 cells while the cells are being washed before fixation. PM: plasma membrane of the Mia PaCa-2 cell. (b) At some point, type I pili can develop binding to mannose residues on the cell surface. This cooperative binding is stronger and resistant to the washing processes. (c) BSUV9c specifically binds to the Mia PaCa-2 cell surface through an unknown mechanism. Because of their motility and affinity, BSUV9c can attach underneath the cell. (d) The BSUV9c on the surface can be detached by washing, but the BSUV9c under or between cells can remain after washing.

As described above for *E. coli* experiments, the same culture condition for Mia PaCa-2 and HPDE cells were tested to confirm that specific binding of mutant *B.subtilis* to Mia PaCa-2 cells is not dependent on the different culture media. Both cells were cultured in Keratinocyte SFM on FN7-10-coated glass surface, and captured movies to see the attachment of bacteria on the cells. As shown in Supplementary movies SI5, 6 the mutant *B.subtilis* specifically bind to the Mia PaCa-2 cells in this condition as well. This suggests that the binding of mutant *B.subtilis* is specific regardless of medium.

### Genomic mutations in BSUV9c

The mutations in the BSUV9c genome were checked by illumina sequencing (Table 2). First, the original lab strain has 26 mutations compared to the 168C sequence in the database (PRJNA57675). Two of them change the amino acid sequences in the genes *uvrX* and *ypqP*. The *uvrX* is reported as a putative UV-damage repair protein in bacteriophage SPβ Although the mutation is a conservative D45E, it might affect survivability after UV treatment. The *ypqP* is an important molecule for biofilm formation in *B. subtilis* isolated from hospitals ^24^. However, since the expression of *ypqP* is disrupted by bacteriophage SPβ in the strain 168C, this mutation probably does not affect results here.

After 9 cycles of mutation/selection processes, 15 new mutations were accumulated in BSUV9c. Three are in genes for membrane proteins (*comP, comQ*, and *yfiZ*), which might affect the cell surface ^25^. Several mutations including *comP, comQ, yjbM, spoIIP*, are in genes known to affect genetic competence and sporulation 26 27 28.

Another possibility is that *fliY* mutation might enhance the polar attachment because a deletion mutant of FliY does not switch direction of motility ^29^. The *fliY* mutations appeared in 10% of the BSUV5 population and reached 100 % in BSUV7 population (see supplementary Fig. S1f online). A potential problem is the mutations in the *mutS* gene, which is a mismatch repair enzyme. If the mutation affects MutS activity, the strain may be genetically unstable.

A clone from BSUV5 (BSUV5c) was investigated as an intermediate strain. This particular clone has a mutation in *rpoB*, *yfiZ*, and *yjbM* (see supplementary Fig. S1 TableS1 online). Because these mutations appeared in ~ 40% of BSUV4 population, which were roughly estimated by the chromatograph for sequencing (see supplementary Fig. S1 online), this BSUV5c probably has a character of BSUV4 mutant. Although 10% of the BSUV5 population (not a cloned strain) already acquired *fliY* mutation and survived to the next cycles, this cloned strain (BSUV5c) did not acquire *fliY* mutation and died out by 7^th^ mutation/selection cycles. The affinity level of BSUV5c to Mia PaCa-2 cells is reasonably intermediate between wt and BSUV9c (Fig. 5 i). The BSUV5c apparently bound 30 times more than to HPDE cells, due to the low affinity of BSUV5c to HPDE cells. Although this is a comparable selectivity to the BSUV9c, the binding to HPDE cells might not be accurate enough because absolute number of bacterial binding to HPDE cells in the experiments is low.

## Discussion

In this study, both Gram-negative *E.coli* and Gram-positive *B.subtilis* increased their affinity to cancer cells through the mutation/selection process. This bacterial evolution occurred in only 10 cycles of the mutation/selection process. Several interesting genes were mutated including those for biofilm-and adhesin-related molecules in *E. coli*. The mutations in *fimB* (the mutation is 3 bp before the transcription start in the *fimB* promoter P1) and *fimE* are probably major factors for the affinity to the Mia Paca-2 cells. FimB and FimE are regulatory recombinases for *fimS*, which is a promoter for type I pili. Both FimB and FimE can recombine and reverse the direction of the *fimS* sequence to suppress the expression of type I pili. This suppression of type I pili is recovered only by the bidirectional recombinase activity of FimE for *fimS*. When both FimB and FimE activities are abolished, *E. coli* should have either 0 or maximum expression of type I pili depending on the fixed direction of fimS. Because type I pili are known to have affinity to mannose residues and adhere to cell surface receptors, the complete inhibition of ECUV10c adhesion to Mia PaCa-2 cells by mannose, a type I pili inhibitor, suggests that ECUV10c upregulates type I pili expression and adheres more strongly to Mia PaCa-2 cells.

In the case of uropathogenic *E. coli*, type I pili adhere to β1/α3 integrin and are involved in the *E. coli* invasion of host epithelial cells. Because Mia PaCa-2 cells express β1/α3 integrin more than normal human pancreatic ductal epithelial cells ^30^, ECUV10c may not only attach more to Mia PaCa-2 cells than HPDE cells but also may be internalized into cells through β1/α3 integrin ^31^.

However, the specificity for Mia PaCa-2 cells is probably not determined by the type I pili expression level alone as described in Results (Fig. 2, 3). One reason is that the ECUV4 population did not have mutations in FimB and FimE genes, but the adhesions to both Mia PaCa-2 and HPDE cells significantly increased compared to the wild type W3110. Another reason is that ECUV10c, which is supposed to upregulate type I pili expression by mutations in FimB and FimE, showed lower adhesion to HPDE cells compared to ECUV4. These facts cannot be explained by type I pili expression level alone. To accommodate these facts, I suggest a two step adhesion process, shown in Fig. 6a, b.

In this model, the molecule(s) that specifically bind to the Mia PaCa-2 cells developed on the ECUV10c surface by the mutation/selection system. At the first step, ECUV10c may weakly but specifically attach to the Mia PaCa-2 cells through this hypothetical molecule(s) (Fig. 6a). Then at the second step, type I pili bind to a cell surface receptor such as β1/α3 integrin Fig. 6b). This binding may also be weak, but the cooperative effect of two weak interactions can generate a very strong binding affinity ^32^. When mannose inhibits the pilus binding it reduces the affinity to the weak binding of the original molecule. The hypothetical molecule(s) which specifically bind to Mia PaCa-2 cells may be related to biofilm formation. This model explains all results obtained here.

In order to kill Mia PaCa-2 cells, *S.aureus* aHL was expressed in ECUV10c from three different pBAD vectors. The expression level of aHL from the original pBAD18 was not enough to kill Mia PaCa-2 cells. Two new vectors, which were modified from pBAD18, could express aHL in ECUV10c at a level that killed Mia PaCa-2 cells. These results are consistent with previous reports that aHL released by *E. coli* after expression from a pBAD vector killed cancer cells. In that study aHL killed breast cancer cells in vitro at the same concentration as in original *E. coli* induction media ^14^. In the present study, Mia PaCa-2 cells were killed at 14 times dilution of the aHL in the original induction media. These results suggest that the combination of strain ECUV10c, pBADMO_aHL and Mia PaCa-2 cells should be optimal not only for specificity to cancer cells, but also aHL activity, compared to the previous report where cancer cells were killed both in cell culture and in mice ^14^.

In the case of *B. subtilis*, BSUV9c showed characteristic adhesions on Mia PaCa-2 cells. When BSUV9c swam and hit the surface of Mia PaCa-2 cells at their poles, they attached for varied periods and some of them stayed for more than 10 min (see Supplementary movieSI 4 online). This distinctive attachment is not strong because the majority of BSUV9c on the top of cells were detached by the washing with PBS before fixation. BSUV9c attached not only on the top of cells but also underneath cells and intercellular space where BSUV9c remained even after washing. The accumulation of BSUV9c underneath cells and intercellular space is not just caused by motile bacteria sticking in narrow spaces because wt strain 168C did not accumulate there.

Although the mechanism by which BSUV9c adheres to Mia PaCa-2 cells is not solved, there are several interesting mutations in the BSUV9c genome. Some mutations may change the motility of *B. subtilis*. A phosphatase FliY regulates the CheY phosphoryration level which determines the rotational direction of flagella ^29^. Because the deletion of FliY is supposed to make *B.subtilis* move forward and reduce the frequency of switching direction, the mutation in the *fliY* gene may inhibit FliY activity and enhance polar attachment by extending the time for pushing the pole to the cells. This may also enhance sticking underneath Mia PaCa-2 cells by unidirectional motility.

Another line of mutations is in genetic competence and sporulation-related genes including *comP*, *comQ*, *yjbM* and *spoIIP*. *YjbM* is one of the ppGpp synthases and affects both competence and sporulation signaling pathways as well as broad cell systems for the stringent responses ^26^. *comP* and *comQ* are mainly involved in the development of genetic competence where comP phosphorylates master regulator *comA* to induce the competence signaling pathway ^27, 28^. However, this pathway also communicates with the sporulation pathway ^27, 28^. One sporulation specific protein, SpoIIP which is a lytic enzyme on the outside of the plasma membrane, might affect the surface of *B. subtilis* ^33, 34^. The sporulation and/or genetic competence genes might be involved in the polar attachment although the expression of sporulation genes under normal growth conditions is not known.

A future step is in vivo experiments using animal models, such as a xenograft mouse model using Mia PaCa-2 cells. It should be tested if accumulation of bacteria on cancer cells can be observed, and can the attached bacteria kill Mia PaCa-2 cells by induction of toxin. ECUV10c is ready to use, since a robust toxin expression system was developed here. The mutation/selection system developed here may be model for bacterial cancer therapy.

**Table 1.** genomic mutations in lab W3110 strain and ECUV10. POS is the position in the W3110 genomic sequence (NCBI Reference Sequence: NC_000964.3). REF is the W3110 sequence. Lab W3110 strain has 9 mutations in the genome, compared to the database W3110 sequence. (a) Bifunctional enzyme 5,10-methylene-tetrahydrofolate dehydrogenase and 5,10-methylene-tetrahydrofolate cyclohydrolase. (b) Predicted enzyme associated with biofilm formation. (c) Predicted diguanylate cyclase. (d) conserved hypothetical protein diguanylate cyclase involed in biofilm formation. (e) RNA polymerase beta prime subunit. (f) carbohydrate-specific outer membrane porin (cryptic). (g) tyrosine recombinase/inversion of on/off regulator of fimA.

| POS | Type | W3110<br>REF | Lab<br>strain<br>w3110_ | ECUV10c | Gene | Protein<br>mutation | Function |
| --- | --- | --- | --- | --- | --- | --- | --- |
| 547694 | SNP | A | G | G | YlbE | E83E | hypothetical protein |
| 547831 | INDEL | A | AG | AG | YlbE | K86Eframeshift | hypothetical protein |
| 556858 | SNP | A | T | T | folD | L36Q | (a) |
| 926691 | SNP | G | REF | A | infA | P59S | translation initiation factor IF-1 |
| 926692 | SNP | G | REF | A | infA | T58T | translation initiation factor IF-1 |
| 928602 | SNP | A | REF | T | cydC | L339Q | ATP binding/permease protein |
| 1065700 | SNP | C | T | T | yccE | R415C | hypothetical protein |
| 1088940 | SNP | C | REF | T | ycdR | G447D | (b) |
| 1093686 | SNP | T | C | C | ycdT | V150A | (c) |
| 1182294 | SNP | A | REF | T | ycfZ | L184STOP | predicted inner memb. preprotein |
| 1197797 | SNP | C | REF | T | icd | H366H | isocitrate dehydrogenase |
| 1197809 | SNP | C | REF | T | icd | T370T | isocitrate dehydrogenase |
| 1197822 | SNP | T | REF | C | icd | L375L | isocitrate dehydrogenase |
| 1197824 | SNP | A | REF | G | icd | L375L | isocitrate dehydrogenase |
| 1197854 | SNP | C | REF | T | icd | N385N | isocitrate dehydrogenase |
| 1197857 | SNP | G | REF | C | icd | A386A | isocitrate dehydrogenase |
| 1197860 | SNP | A | REF | G | icd | K387K | isocitrate dehydrogenase |
| 1197869 | SNP | C | REF | T | icd | T390T | isocitrate dehydrogenase |
| 1197881 | SNP | G | REF | A | icd | E394E | isocitrate dehydrogenase |
| 1303712 | SNP | A | G | G | oppA | S273G | oligopeptide transporter |
| 1317208 | SNP | C | REF | T | yciF | E93K | hypothetical protein |
| 1624961 | SNP | C | REF | A | ydeH | V202F | (d) |
| 1646060 | SNP | G | REF | A | intergenic |  |  |
| 2019202 | SNP | C | T | T | flil | T171I | flagellum ATP synthase |
| 2627746 | SNP | C | REF | T | intergenic |  |  |
| 2906975 | SNP | C | G | G | pyrG | D450H | CTP synthetase |
| 3449836 | SNP | A | REF | C | rpoC | I499S | (e) |
| 3735150 | SNP | G | REF | A | bglH | V230V | (f) |
| 3735183 | SNP | G | REF | T | bglH | Q241H | (f) |
| 4371270 | INDEL | GAA | G | G | intergenic |  |  |
| 4421323 | SNP | T | REF | A | yjfO | K25STOP | hypothetical memb. protein |
| 4542607 | INDEL | GC | REF | G | yjhT | A277frameshift | hypothetical protein |
| 4545485 | SNP | G | REF | T | intergenic<br>(fimB) |  | fimB promoter P1 |
| 4547053 | SNP | C | REF | T | fimE | R115C | (g) |

**Table 2.**
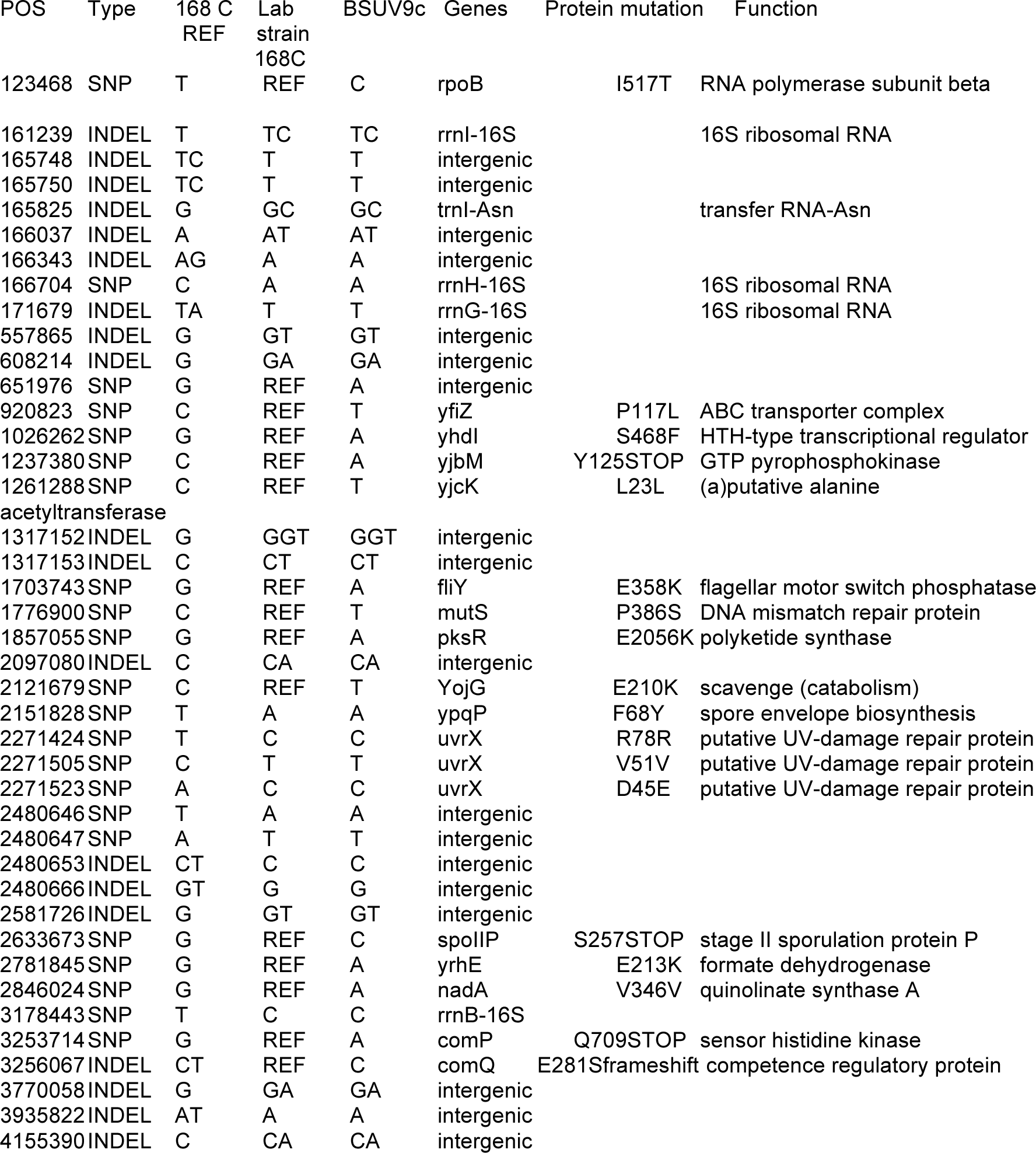
genomic mutations in lab *B. subtilis* 168C strain and BSUV9. POS is the position for 168C genomic sequence in FASTA (PRJNA57675). REF is the *B. subtilis* 168C sequence. Lab *B. subtilis* 168C strain has 26 mutations in the genome, compared to the *B. subtilis* 168C sequence in FASTA.

## Acknowledgements

I thank Dr. Rebekah. R White and Dr. Partha. Ray for extensive discussion and supporting cell culture. I thank Dr. Harold P. Erickson for critical reading of manuscript. This research was supported by National Institutes of Health Grant CA 047056 to Dr. Harold P. Erickson.

